# Ketogenic diet prevents obesity-associated pancreatic cancer independent of weight loss and induces pancreatic metabolic reprogramming

**DOI:** 10.1101/2025.05.20.655200

**Authors:** Ericka Vélez-Bonet, Kristyn Gumpper-Fedus, Kaylin Chasser, Zachary Hurst, Hsiang-Yin Hsueh, Valentina Pita-Grisanti, Alexus Liette, Grace Vulic, Fouad Choueiry, Huan Zhang, Jiangjiang Zhu, Sue E. Knoblaugh, Stacey Culp, Jeff S. Volek, Zobeida Cruz-Monserrate

**Author notes:** Correspondence (Z.C-M.).

## Abstract

Pancreatic ductal adenocarcinoma (PDAC) is an aggressive cancer with poor outcomes. Obesity is a risk factor for several cancers including PDAC due to metabolic dysregulation and inflammation. The ketogenic diet (KD) can alter metabolism and has been evaluated for its effects on tumor progression in non-obese but not obese PDAC using genetically engineered mouse models (GEMMs). We hypothesized that ketone bodies and a KD alter cell and tumor metabolism. We show that ketone treatments altered pyrimidine metabolism in PDAC cells. Moreover, in an obese PDAC GEMM, KD prevented tumor progression independent of weight loss but promoted PDAC in a non-obese PDAC GEMM. The KD-specific delay of obesity-associated PDAC was associated with pancreatic metabolic shifts in pyrimidine, cysteine and methionine, and arginine and proline pathways. These findings suggest potential benefits of a KD in preventing obesity-associated PDAC, but highlights some risks in non-obese settings.

## Introduction

Pancreatic ductal adenocarcinoma (PDAC) is one of the most aggressive and lethal cancers, with a five-year survival rate of 13%, largely due to late diagnosis and resistance to conventional therapies.^1^ Addressing these challenges requires advancing preventive measures and therapeutic strategies. Obesity, a well-established risk factor for PDAC, contributes to tumor progression through chronic inflammation, insulin resistance, and altered adipokine signaling, which promotes a pro-tumorigenic microenvironment.^2–4^ Furthermore, obesity driven metabolic dysregulation alters key pathways involved in energy balance and systemic metabolism.^4^ This leads to metabolic adaptations in nutrient limited environments that support cancer cell survival,^5^ highlighting a need to understand how metabolic interventions may modulate tumor growth.^2^ Epidemiological data indicate that obesity prevalence continues to rise globally, driven by dietary patterns that contain high amounts of sugar, red meat, and processed foods, typically characterized as “Western” diets, and sedentary lifestyles.^6,7^ Obesity management through lifestyle interventions, such as dietary modifications and physical activity, could reduce cancer risk.^8–11^ Given that obesity driven metabolic reprogramming influences tumor progression, understanding how tumors adapt their metabolism is critical for developing targeted preventive strategies for obesity-associated PDAC.

Metabolic reprogramming is a hallmark of many cancers, including PDAC, where cells have an increased reliance on glucose metabolism and biosynthetic pathways that sustain rapid proliferation.^12–14^ As a result, PDAC cells adapt to fluctuations in nutrient availability, particularly through increased glucose uptake and glycolytic flux.^15^ Cancer cells rely on aerobic glycolysis to convert glucose to lactate, known as the Warburg effect, to sustain growth.^16^ The ketogenic diet (KD), which elevates ketone bodies in circulation, has been studied for its effect on metabolic disorders, such as obesity and cancer.^17–20^ They can promote weight loss and improve glycemic control by lowering circulating glucose and insulin, even in stage IV cancers.^17–20^ Short term (≤ 8 weeks) consumption of a KD reduces inflammatory markers and this effect is enhanced in obese individuals.^21^ However, long term consumption of a KD in a non-obese mouse can cause glucose intolerance, inflammation, and temporary weight loss.^22^ Non-obese PDAC genetically engineered mouse models (GEMMs) fed a KD still have tumor development; however, KDs enhance therapeutic responses when combined with agents such as gemcitabine, nab-paclitaxel or cisplatin, ultimately improving survival outcomes.^23–25^ Therefore, while KDs have demonstrated metabolic benefits in obesity, improving glucose regulation and modifying systemic metabolism,^26–28^ their potential to prevent tumor initiation and progression in obesity-associated PDAC is unclear and worth studying.

This study investigates differences in ketone body treatments on PDAC cell metabolism and the effects of a KD on tumor development in obese and non-obese preclinical PDAC models. Moreover, we examined pancreas-specific metabolomic differences associated with the dietary modifications utilized after obesity-associated PDAC. Our goal was to test whether a KD intervention could prevent obesity-associated PDAC and study the associated pancreas-specific metabolic reprograming that could be leveraged for tumor prevention.

## Results

### Ketone bodies promote pyrimidine metabolism in PDAC cells

To test whether ketone bodies modulate key metabolic pathways in PDAC cells, we treated LSL-Kras^G12D^/LSL-Trp53^-/-^/Pdx1-CRE (KPC) cells^29^ with sodium 3-hydroxybutyrate (NaHB) or lithium acetoacetate (LiAcAc), salt forms of the two most common ketone bodies in circulation, beta-hydroxybutyrate (β-HB) and acetoacetate (AcAc),^30,31^ and performed untargeted metabolomic analysis (**Figure 1A**). Partial least square discriminant analysis (PLS-DA) showed that the NaHB, LiAcAc, and control treated PDAC cells clustered separately, indicating unique differences in metabolite composition based on treatment (**Figure 1B**). A metabolite set enrichment analysis (MSEA) using the Kyoto Encyclopedia of Genes and Genomes (KEGG) database identified the enriched pathways within PDAC cells after treatment with NaHB (**Figure 1C**) or LiAcAc (**Figure 1D**) compared to control treated PDAC cells, respectively. Of these, only 10 pathways were significantly enriched in the NaHB treated cells while only 6 pathways were significantly enriched in the LiAcAc treated cells. Across the treatment groups, 35 metabolites had significant differential abundance highlighting key metabolic changes (**Figure 1E**).

**Figure 1:**
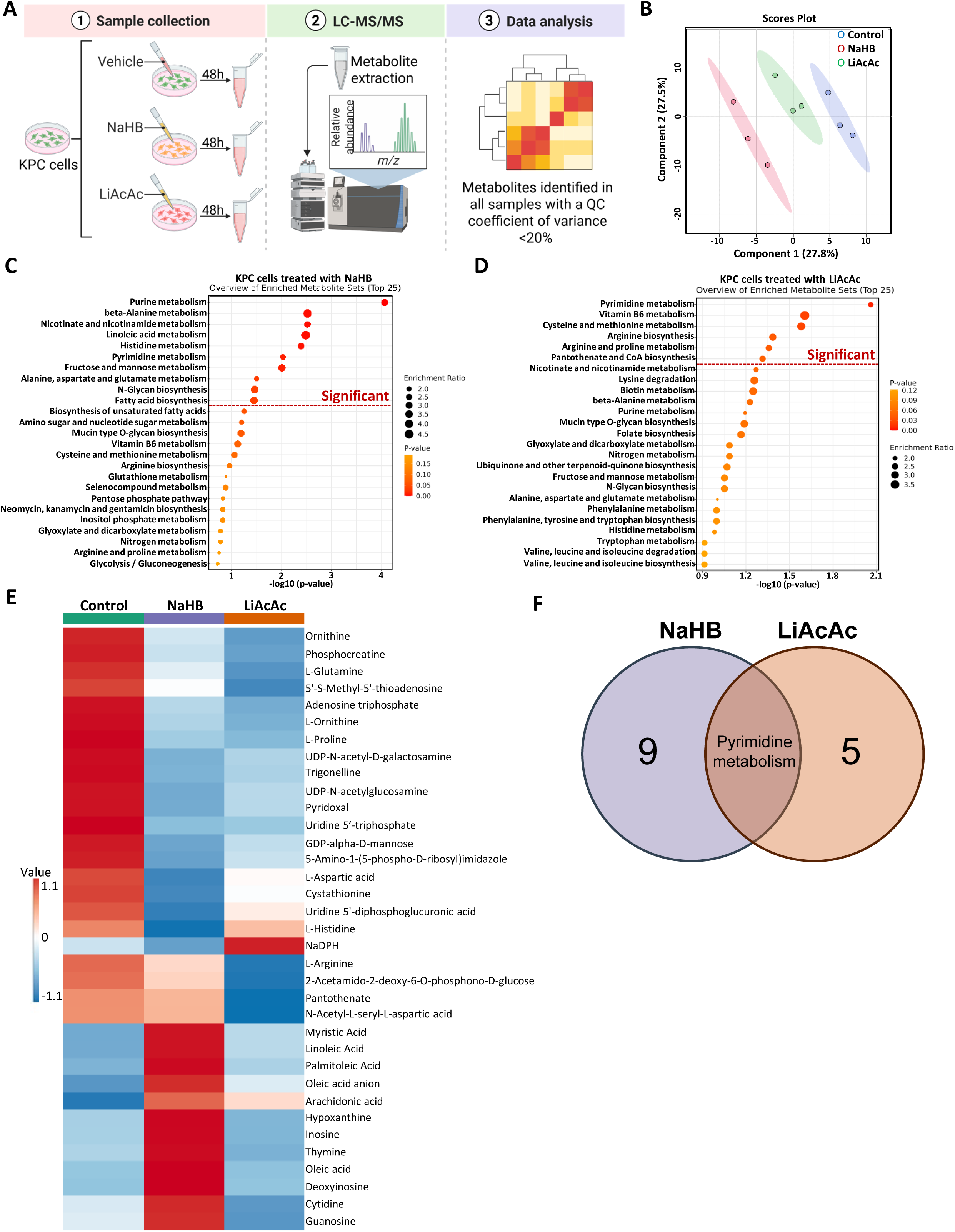
Metabolomic profiling of PDAC cells identifies pyrimidine metabolism as a common pathway altered in response to NaHB and LiAcAc treatments. **A)** Workflow for untargeted metabolomics of KPC cells treated with ketone bodies. **B**) PLS-DA plot showing clustering between KPC cells treated with ketone bodies or a vehicle control (n=3). KEGG pathways analysis when KPC cells are treated with **C**) NaHB or **D**) LiAcAc. **E**) Heatmap summarizing the relative abundance of metabolites with p<0.05. **F**) Venn diagram of the significantly altered pathways in cells treated with ketone bodies.

Pyrimidine metabolism emerged as the common pathway influenced by both ketone body treatments (**Figure 1F**). Notably, NaHB treatment enriched pyrimidine metabolism by increasing cytidine and thymine and decreasing uridine 5’-triphosphate (UTP) and glutamine abundance.

Similarly, LiAcAc treated cells also downregulated UTP and glutamine. Additionally, LiAcAc treatment affected arginine and proline metabolism by decreasing L-proline, L-arginine, phosphocreatine and L-ornithine abundance compared to vehicle control treated cells. These data highlight the specific metabolic alterations ketone bodies cause to PDAC cells.

### KD delays tumor development in obesity-associated PDAC GEMMs independent of weight loss

To determine whether a KD could prevent obesity-associated PDAC tumor development, obesity was induced in control and LSL-Kras^G12D^/Elastase-Cre^ERT^ (Kras^G12D^) mice.^32^ The mice were subjected to diet-induced obesity (DIO) using a 45% high-fat diet (HFD), as it mimics a more physiologically relevant DIO in humans,^33^ for 15 weeks. Mice were then randomized to be maintained on the DIO or switched to either a KD or a low-fat control to the KD (KDC) for 6 weeks (**Figure 2A**).

**Figure 2:**
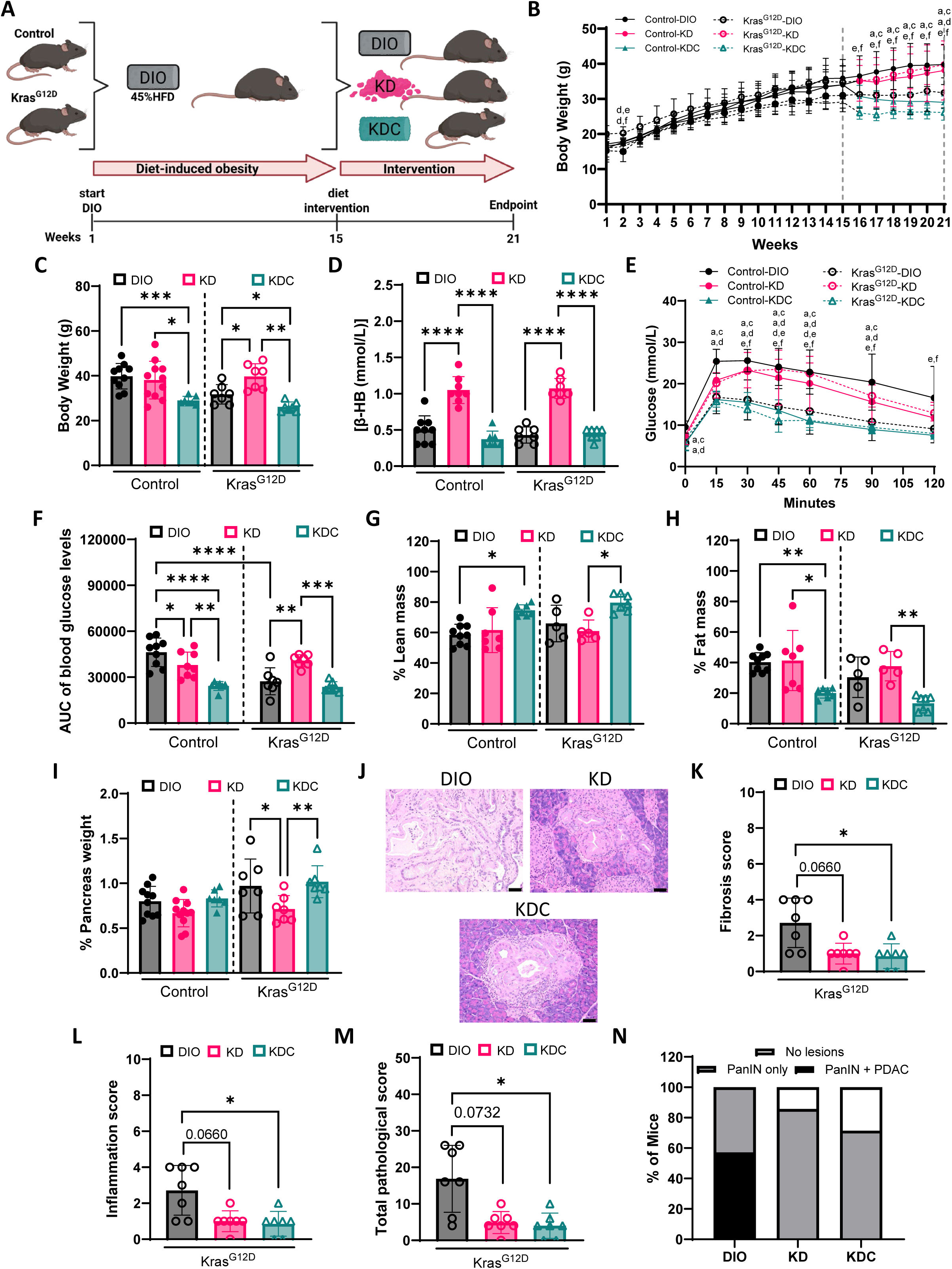
KD delays tumor development in an obesity-associated PDAC GEMM independent of weight loss using a DIO that is physiologically relevant to humans. **A)** Experimental design to evaluate the effects of the KD and KDC following diet-induced obesity (DIO) using 45% high-fat diet (HFD) in an obese PDAC GEMM. **B**) Body weight of control and Kras^G12D^ mice over 21 weeks (control n=28; Kras^G12D^ n=21). **C**) Body weight at the study endpoint (week 21) following dietary interventions (n=7-11 mice/group). **D**) β-OH concentration over the 6-week intervention in control and Kras^G12D^ mice. GTT measuring **E**) glucose and **F**) area under the glucose curve (AUC) at endpoint. Percent **G**) lean mass, **H**) fat mass, and **I**) pancreas weight of control and Kras^G12D^ mice at endpoint. **J**) Representative hematoxylin and eosin (H&E) stains of pancreas sections from Kras^G12D^ mice. Original magnification, ×20. Scale bar, 50 µm (black bar). Pathology scores for **K**) fibrosis, **L**) inflammation, and **M**) total pathological score (n=7 mice/group). **N**) Percent of Kras^G12D^ mice that developed PDAC or mouse pancreatic intraepithelial neoplasia (mPanIN) lesions. Statistical significance determined by mixed-between-within ANOVA’s where significance between groups indicated by letter pairs listed for control a) DIO b) KD and c) KDC and Kras^G12D^ d) DIO, e) KD and f) KDC (**B** and **E**), one-way ANOVA with Brown Forsythe and Dunnet’s corrections (**C**), one-way ANOVA with Tukey’s correction for multiple comparisons (**D**), two-way ANOVA with Tukey’s correction for multiple testing (**F**), Kruskal-Wallis test with Dunn’s correction for multiple testing (**G**, **H**, and **K-M**), or one-way ANOVA with Sídak correction for multiple comparisons (**I**), *p<0.05, **p<0.01, ***p<0.001, ****p<0.0001

After dietary interventions, control KDC mice had lower body weight compared to control DIO and KD mice (**Figure 2B and C**). Kras^G12D^ KD mice had higher body weight than those on DIO and on a KDC diet intervention. Furthermore, Kras^G12D^ KDC mice had lower body weight than Kras^G12D^ DIO mice.

β-HB concentration was elevated in both control and Kras^G12D^ KD mice compared to KDC and DIO mice (**Figure 2D**). In control mice, a glucose tolerance test (GTT) showed that DIO and KD mice had impaired glucose tolerance compared to KDC mice (**Figure 2E and F**). Furthermore, control DIO mice had impaired glucose tolerance compared to control KD mice and Kras^G12D^ DIO mice. Additionally, Kras^G12D^ KD mice had impaired glucose tolerance compared to Kras^G12D^ DIO and KDC mice. There was an increase in percent lean mass in control KDC mice compared to control DIO mice and Kras^G12D^ KDC mice compared to the Kras^G12D^ KD mice (**Figure 2G**). Control KDC mice had lower percent fat mass compared to control DIO and KD mice (**Figure 2H**). In addition, Kras^G12D^ KDC mice had a lower percent fat mass than Kras^G12D^ KD mice. Pancreas weight as a percent of the body weight was lower in Kras^G12D^ KD mice compared to Kras^G12D^ DIO and KDC mice (**Figure 2I**). Kras^G12D^ KDC and KD mice had less pancreatic fibrosis (**Figure 2J and K**), inflammation (**Figure 2L**), and overall pathological scores (**Figure 2M**) compared to DIO mice. Notably, PDAC developed in 57% of the Kras^G12D^ DIO mice, whereas no PDAC was observed in the Kras^G12D^ KD or KDC mice (**Figure 2N**).

Instead, KD and KDC mice only had mouse pancreatic intraepithelial neoplasia (mPanIN) lesions or no lesions. Overall, these results suggest that the KD delays tumor development in obesity-associated PDAC, like KDC, but relies on a mechanism independent of weight loss.

We conducted another experiment to determine whether a DIO higher in fat content (60% HFD, commonly used in mouse obesity and cancer studies)^11,32^ results in similar outcomes as a KD intervention after the more physiologically relevant 45% DIO (**Figure 3A**). Following DIO, Kras^G12D^ on the KD intervention had higher body weight than control KD mice and Kras^G12D^ KDC mice (**Figure 3B and C**). β-HB concentration was elevated in control KD mice compared to control KDC mice and Kras^G12D^ KD mice compared to Kras^G12D^ 60% DIO mice (**Figure 3D**). In control mice, a GTT showed that DIO and KD mice had impaired glucose tolerance compared to KDC mice. (**Figure 3E and F**). In addition, Kras^G12D^ DIO mice had impaired glucose tolerance compared to KDC mice; however, the KD did not worsen glucose tolerance compared to a 60% DIO. Control KD or KDC mice had more lean mass and less fat mass compared to control 60% DIO mice while Kras^G12D^ KDC mice had more lean mass and less fat mass compared to KD and 60% DIO mice (**Figure 3G and H**). The pancreas weight as a percent of the body weight was increased in Kras^G12D^ KDC mice compared to KD and 60% DIO mice (**Figure 3I**). Fibrosis scores remained unchanged, but less inflammation was observed in the KD mice compared to 60% DIO mice (**Figure 3J-L**). Furthermore, the total pathological score was significantly lower in Kras^G12D^ KD mice compared to Kras^G12D^ 60% DIO mice (**Figure 3M**). In this model, some KDC mice did develop PDAC; however, none of the KD mice did (**Figure 3N**). In contrast, consuming a KD did not affect tumor growth in an obese orthotopic model of PDAC (**Supplemental Figure 1**). These findings suggest that the KD can also delay tumor development following a DIO higher in fat content, independent of weight loss.

**Figure 3:**
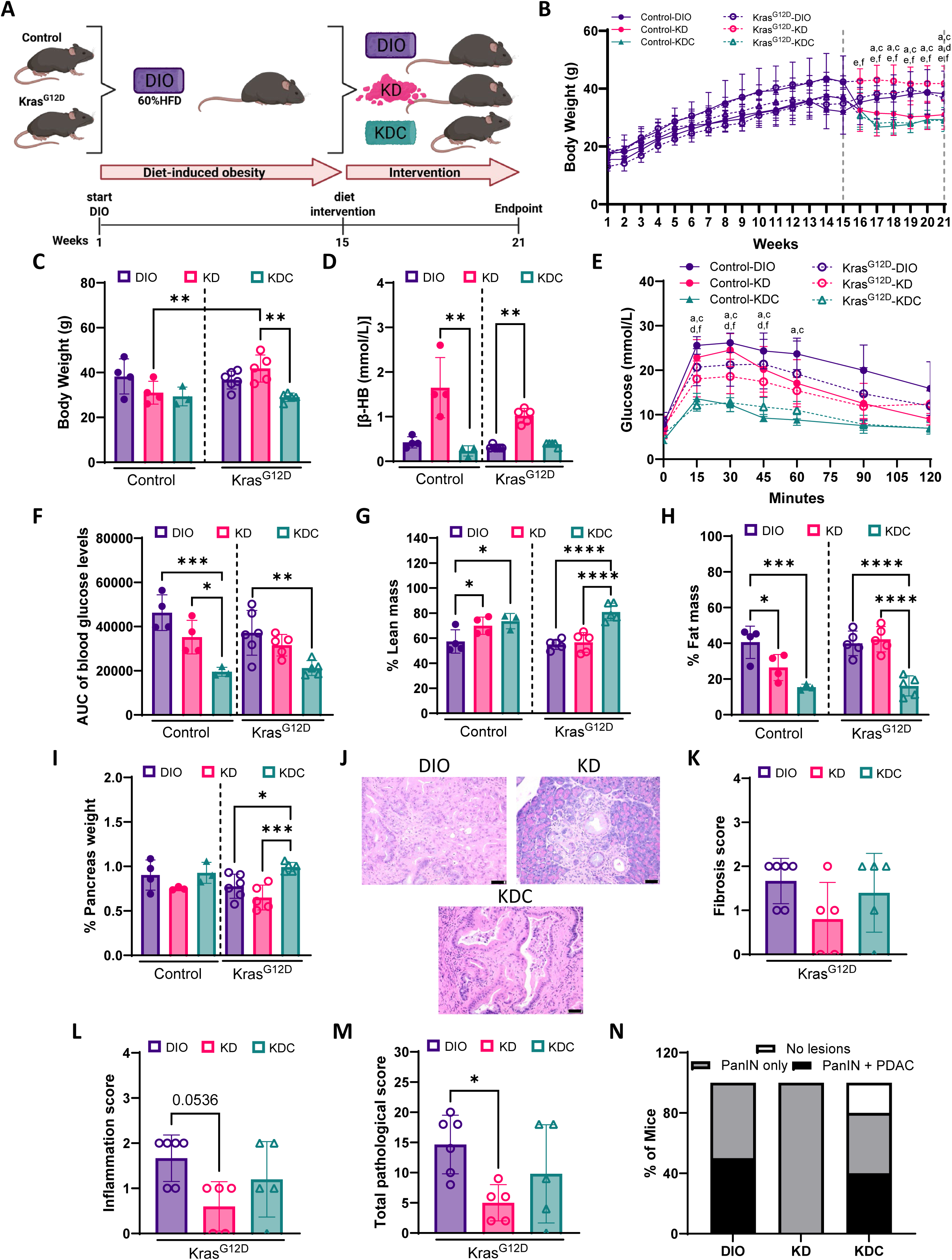
KD delays tumor development in an obesity-associated PDAC GEMM independent of weight loss using a DIO higher in fat content. **A)** Experimental design to evaluate the effects of the KD and KDC following DIO using a 60% HFD in an obese PDAC GEMM. **B)** Body weight of control and Kras^G12D^ mice over 21 weeks (control n=11; Kras^G12D^ n=16). **C**) Body weight at the study endpoint (week 21) following dietary interventions (n=3-6 mice/group). **D**) β-OH concentration over the 6-week intervention in control and Kras^G12D^ mice. GTT measuring **E**) glucose and **F**) AUC at endpoint. Percent **G**) lean mass, **H**) fat mass, and **I**) pancreas weight of control and Kras^G12D^ mice at endpoint. **J**) Representative H&E stains of pancreas sections of Kras^G12D^ mice. Original magnification, ×20. Scale bar, 50 µm (black bar). Pathology scores for **K**) fibrosis, **L**) inflammation, and **M**) total pathological score (n=5-6 mice/group. **N**) Percent of Kras^G12D^ mice that developed PDAC or mPanIN lesions. Statistical significance determined by mixed-between-within ANOVA’s where significance between groups indicated by letter pairs listed for control a) DIO, b) KD, and c) KDC and Kras^G12D^ d) DIO, e) KD and f) KDC (**B** and **E**), two-way ANOVA with Tukey’s correction for multiple testing (**C**, **G**, and **H**), Kruskal-Wallis test with Dunn’s correction for multiple testing (**D** and **K-M**), or one-way ANOVA with Sídak correction for multiple comparisons (**F** and **I**) *p<0.05, **p<0.01, ***p<0.001, ****p<0.0001

### In the non-obese setting, KD promotes PDAC development

To determine whether a KD would prevent PDAC in the non-obese setting, we fed control and Kras^G12D^ mice a low-fat (non-obese) diet for 15 weeks (**Figure 4A**). Mice were randomized to be maintained on the non-obese diet or switched to either a KD or a KDC for a 6-week intervention concluding on week 21.

**Figure 4:**
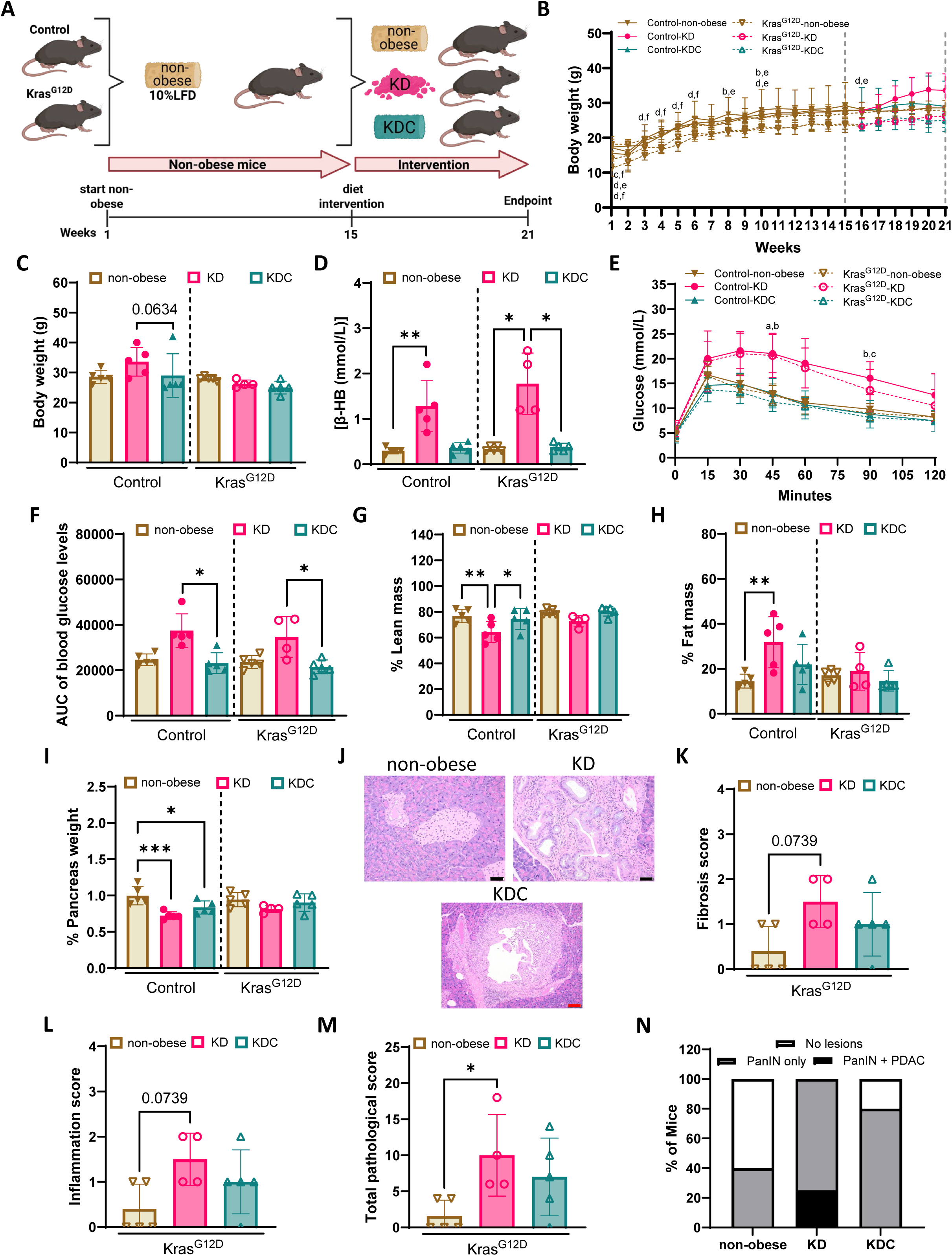
KD promotes tumor development in a non-obese PDAC GEMM. **A)** Experimental design to evaluate the effects of KD, and KDC in a lean PDAC GEMM. **B**) Body weight of control and Kras^G12D^ mice over 21 weeks (control n=15; Kras^G12D^ n=14). **C**) Body weight at the study endpoint (week 21) following dietary interventions (n=4-5 mice/group). **D**) β-OH concentration over the 6-week intervention in control and Kras^G12D^ mice. GTT measuring **E**) glucose and **F**) AUC at endpoint. Percent **G**) lean mass, **H**) fat mass, and **I**) pancreas weight of control and Kras^G12D^ mice at endpoint. **J**) Representative H&E stains of pancreas sections of Kras^G12D^ mice. Original magnification, x20 and x10. Scale bar, 50 µm (black bar) and 100 µm (red bar). Pathology scores for **K**) fibrosis, **L**) inflammation, and **M**) total pathological score (n=4-5 mice/group). **N**) Percent of Kras^G12D^ mice that developed PDAC or mPanIN lesions. Statistical significance determined by mixed-between-within ANOVA’s where significance between groups indicated by letter pairs listed for control a) non-obese, b) KD and c) KDC and Kras^G12D^ d) non-obese, e) KD and f) KDC (**B** and **E**), Kruskal-Wallis test with Dunn’s correction for multiple testing (**C, D, F**, and **K-M**), or one-way ANOVA with Sídak correction for multiple comparisons (**G, H**, and **I**). *p<0.05, **p<0.01, ***p<0.001, ****p<0.0001.

Following the 6-week intervention, control KD mice were slightly heavier than the control KDC mice (**Figure 4B and C**). β-HB concentration was elevated in control KD mice compared to non-obese mice and in Kras^G12D^ KD mice compared to non-obese and KDC mice (**Figure 4D**). GTT showed impaired glucose tolerance in control and Kras^G12D^ KD mice compared to KDC mice (**Figure 4E and F**). Control KD mice had less lean mass compared to non-obese and KDC mice, and more fat mass compared to non-obese controls (**Figure 4G and H**). In contrast, Kras^G12D^ mice had no differences in lean and fat mass across the diet groups.

Furthermore, pancreas weight as a percent of the body weight was lower in control KD and KDC mice compared to non-obese mice; however, there was no difference in percent pancreas weight in the Kras^G12D^ mice across diet groups (**Figure 4I**). Kras^G12D^ KD mice had an increased trend in pancreatic fibrosis and inflammation, along with a significant increase in overall pathological score compared to Kras^G12D^ non-obese mice (**Figure 4J-M**). Notably, 25% of the KD mice developed PDAC (**Figure 4N**). Similarly, consuming a KD did not affect tumor growth in a non-obese orthotopic model (**Supplemental Figure 2**). These data demonstrated that in a non-obese setting, KD intervention did not delay tumor development, rather it promoted PDAC.

### KD induces tumor metabolic reprogramming in obesity-associated PDAC

Given that the PDAC GEMMs with DIO fed a KD did not develop PDAC (**Figures 2 and 3**) and ketones induced metabolic reprogramming of PDAC cells (**Figure 1**), we examined the effects of the KD on the metabolomic profiles of the pancreatic tissue of mice which were fed the more physiologically human relevant DIO (45% HFD, **Figure 2**).^33^ A PLS-DA showed distinct metabolic clustering between DIO, KD, and KDC (**Figure 5A**). Enrichment analysis based on the KEGG database identified the enriched metabolic pathways between KD and DIO (**Figure 5B**) and between KDC and DIO (**Figure 5C**). Of the top 25 enriched metabolite sets identified, 9 were significantly enriched when comparing Kras^G12D^ KD mice to Kras^G12D^ DIO mice and 8 were significantly enriched when comparing Kras^G12D^ KDC mice to Kras^G12D^ DIO mice. These pathways were driven by 56 differentially abundant metabolites between the KD and DIO mice and 51 differently abundant metabolites between the KDC and DIO mice (**Figure 5D and E**). Between the KD and DIO mice and the KDC and DIO mice, there were 3 common pathways significantly enriched which included, pyrimidine metabolism, consistent with the pathway identified in PDAC cells treated with ketone bodies. In contrast, arginine and proline and cysteine and methionine metabolism, were significantly enriched only in Kras^G12D^ KD mice compared to Kras^G12D^ DIO mice, but not in Kras^G12D^ KDC mice, which lost weight. Interestingly, arginine and proline metabolism and cysteine and methionine metabolism were also enriched in PDAC cells treated with LiAcAc. These findings indicate that the delay in tumor development due to the KD, independent of weight loss, was associated with changes in pancreas metabolism.

**Figure 5:**
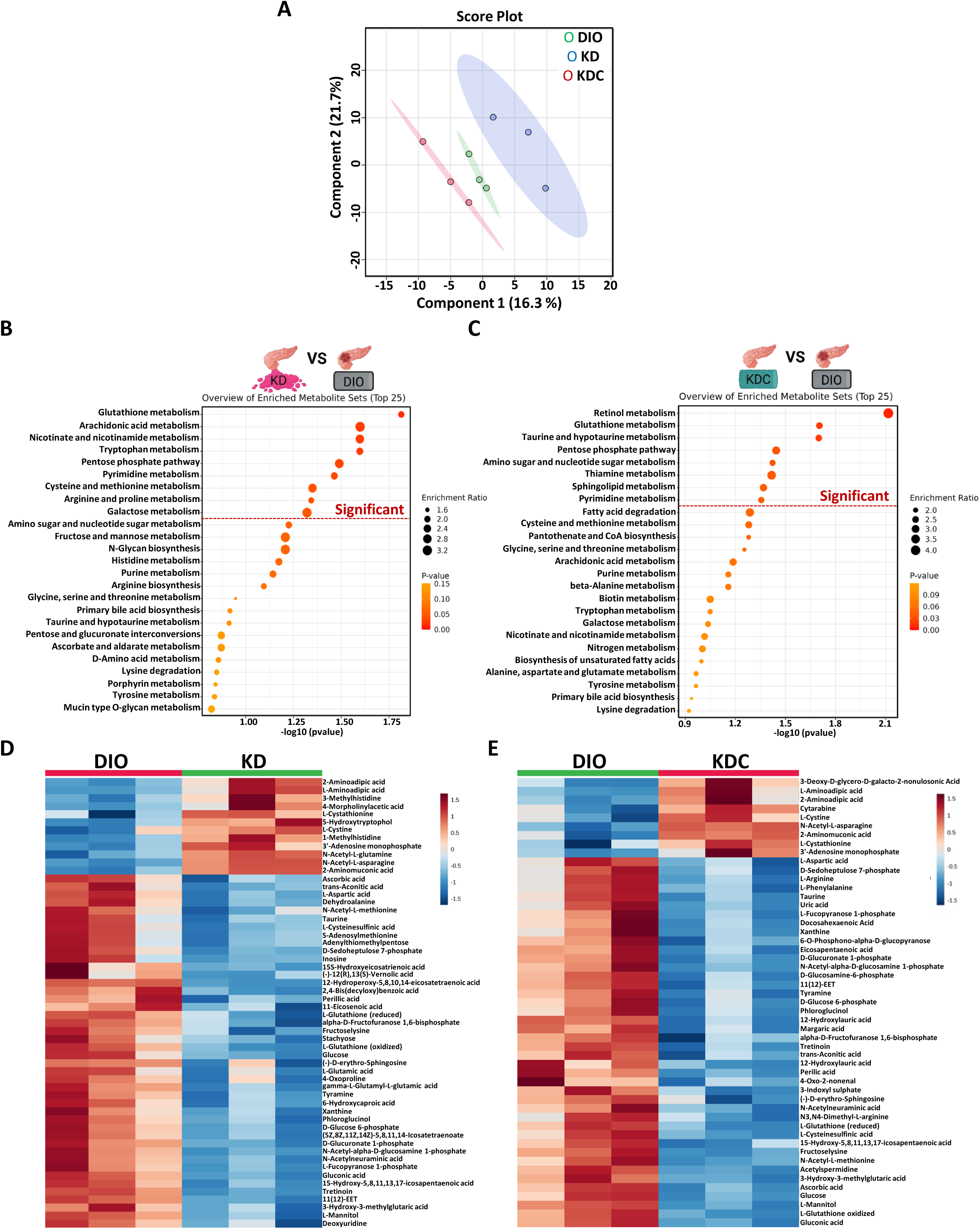
Metabolic profiling of pancreas tissues in an obese PDAC GEMM following dietary interventions. **A)** PLS-DA plot depicting distinct metabolic profiles of each dietary intervention in an obese PDAC GEMM (n=3). KEGG pathway analysis of metabolic pathways in pancreas samples comparing **B**) KD to DIO mice or **C**) KDC to DIO mice. **D**) Heatmap of relative abundance of altered metabolites with p<0.05 between pancreas samples on DIO compared to KD and **E**) DIO compared to KDC.

## Discussion

Since obesity is a risk factor for PDAC,^2,3^ obesity management strategies involving lifestyle interventions like diet and/or physical activity modifications have been investigated for obesity-associated cancer prevention.^9,11,34^ In this study, we showed that consumption of a KD after DIO can delay obesity-associated PDAC development independent of weight loss in a PDAC GEMM. However, a KD promoted PDAC development in a non-obese model.

Furthermore, an untargeted metabolomic approach in both *in vitro* and *in vivo* models of PDAC suggest that ketones or a KD induces metabolic changes that may contribute to preventing tumor development in an obese PDAC GEMM.

Until now, cancer studies on the KD have used non-obese models or have not induced obesity prior to using a KD to assess the effectiveness of the KD at treating or preventing cancer.^23,24,35,36^ Although, KDs can promote weight loss, our findings indicate that PDAC GEMMs that were previously obese and switched to a KD had reduced cancer incidence without concurrent weight loss, suggesting a cancer prevention mechanism that is independent of weight loss. Furthermore, although a KDC intervention that causes weight loss was sufficient to delay tumor development after a relevant physiological form of DIO, it was insufficient to delay tumor development after a higher-fat form of DIO. This was likely due to the higher-fat DIO’s stronger induction of pancreatic inflammation and neoplastic reprogramming that persists despite subsequent diet-induced weight loss.^11,32,37^ Together, this suggests that prevention of obesity-associated PDAC may require a form of metabolic rewiring that is not fully achieved with diet-induced weight loss. It is not without precedent that the benefits of a KD can be observed in absence of weight loss, such as in a model of type 2 diabetes.^38^ While a KD can reduce inflammation in cancer,^39^ our findings show that the KD did not significantly suppress PDAC development in a non-obese GEMM, suggesting that the potential cancer prevention benefits of the KD may be specific to obese individuals.

Obesity is associated with impaired glucose homeostasis due to insulin resistance which can cause hyperinsulinemia, driving PDAC tumorigenesis.^37^ The KD normally influences metabolism lowering circulating glucose and insulin to regulate some of the metabolic dysfunction associated with obesity.^31,40,41^ In this study, non-cancer mice switched to the KD after DIO had improved glucose tolerance, consistent with the known benefits of consuming a KD.^42^ However, in this study, mice carrying pancreatic-specific mutant Kras^G12D^ had impaired glucose tolerance after switching to a KD compared to mice switched to the KDC, a phenotype similar to glucose intolerance that develops after a long-term KD in non-obese mouse models.^22,42^ Interestingly, we show Kras^G12D^ KD mice after DIO did not have histological evidence of tumors even though the transition of PanIN to PDAC is promoted by high levels of glucose and insulin receptor signaling in obese PDAC GEMMs.^37^ Our data suggest that in obese mice with pancreatic-specific mutant Kras^G12D^, the tumor prevention effect of the KD is also independent of managing glucose handling and is likely through other metabolic pathways.

The KD increases circulating ketone bodies, such as β-HB and AcAc, which have been investigated for their metabolic and signaling roles in cancer.^31,43–45^ Here, β-HB was elevated in the obese and non-obese PDAC GEMMs after KD intervention. However, in other non-obese PDAC models, β-HB can promote tumor growth and progression to support fatty acid synthesis under lipid-limited conditions.^46–48^ This aligns with our data showing non-obese PDAC GEMMs had increased pancreatic fibrosis, inflammation, and a higher incidence of PDAC. Since the obese PDAC GEMM had increased β-HB after KD intervention and no tumor development, elevated circulating β-HB may be suppressing tumor development when lipid supply is sufficient but promoting tumor development when lipid availability is restricted. Ketone body supplementation reduces tumor cell viability,^49^ supporting the possibility that β-HB impairs tumor progression. Together, these findings suggest that increasing circulating β-HB through a KD or ketone body supplementation could offer a targeted strategy to delay obesity-associated PDAC development.

Untargeted metabolomics in both *in vitro* and *in vivo* models of PDAC identified pyrimidine, cysteine and methionine, and arginine and proline metabolic pathways as enriched due to ketone body or KD intervention. When we treated PDAC cells with NaHB and LiAcAc pyrimidine metabolism was enriched, which suggests a shift in nucleotide biosynthesis that could influence tumor cell proliferation.^50,51^ In the obesity-associated PDAC GEMM, the pyrimidine pathway was also enriched in Kras^G12D^ KD and KDC mice compared to DIO, but the downregulated metabolites were different, suggesting a selective modulation of nucleotide metabolism specific to the KD after DIO that may inhibit tumor development.^52^ Interestingly, tumor-associated macrophages in PDAC secrete pyrimidine species that inhibit the effectiveness of chemotherapies like gemcitabine.^50^ Since the KD downregulated these metabolites in our pancreatic tissue metabolomics data, this may explain why the KD synergizes with anti-cancer therapies in other models of cancer in the non-obese setting.^23–25^

The cysteine and methionine pathway was significantly enriched in Kras^G12D^ KD mice compared to DIO and in PDAC cells treated with LiAcAc. Changes in this pathway may be strengthening antioxidant defenses and nutrient biosynthesis in the pancreas under metabolic stress.^53,54^ Since decreasing methionine metabolism represents a therapeutic dependency in cancer,^55–57^ such metabolic shifts may be related to the reduced fibrosis, inflammation, and tumor development observed in the Kras^G12D^ KD mice after DIO.

A metabolic pathway that was enriched in Kras^G12D^ KD mice compared to DIO, but not the KDC mice compared to DIO, was the arginine and proline metabolism pathway. This was also enriched in PDAC cells treated with LiAcAc. Arginine metabolism contributes to collagen production through the synthesis of proline to support extracellular matrix remodeling and angiogenesis, key features of the tumor microenvironment.^58^ Here, the metabolites of arginine and proline metabolism were downregulated in PDAC cells treated with LiAcAc and in Kras^G12D^ KD mice which had reduced fibrosis and inflammation. This suggests that ketone bodies and the KD may influence stromal remodeling in a way that suppresses desmoplasia in obesity-associated PDAC. Together, these findings suggest that while some metabolic pathways are common to both dietary interventions, ketone body- or KD-driven alterations may uniquely contribute to tumor-suppressive metabolic reprogramming in obesity-associated PDAC independent of weight loss.

Overall, we demonstrate that a KD may modify circulating nutrient availability to prevent tumor development in obesity-associated PDAC models, but not in non-obese PDAC models.

Furthermore, the pancreatic enriched metabolic pathways specific to KD after DIO are potential intervention targets for delaying PDAC development in high-risk individuals with obesity, independent of weight loss.

### Limitations of the study

This study establishes a foundation for understanding KD induced metabolic adaptations in obesity-associated PDAC as a possible preventive strategy in high-risk individuals with obesity. This study has some limitations, such as the exclusive use of male mice, which may not fully capture the sex-specific responses to the KD after DIO. Additionally, while untargeted metabolomics provided valuable insights into the KD induced tumor metabolic shifts, it did not identify the upstream drivers of these changes. This approach identifies enriched metabolic pathways but does not distinguish whether they result from direct metabolic reprogramming or secondary adaptations to metabolic stress. Integrating transcriptomic, proteomic, and metabolic flux analyses may further clarify whether these metabolic shifts were driven by gene expression changes or enzyme activity which are subject to future studies.

## STAR Methods

### Lead contact

Further information and requests for resources and reagents should be directed to and will be fulfilled by the Lead Contact, Zobeida Cruz-Monserrate

### Materials availability

This study did not generate new unique reagents.

### Key resources table

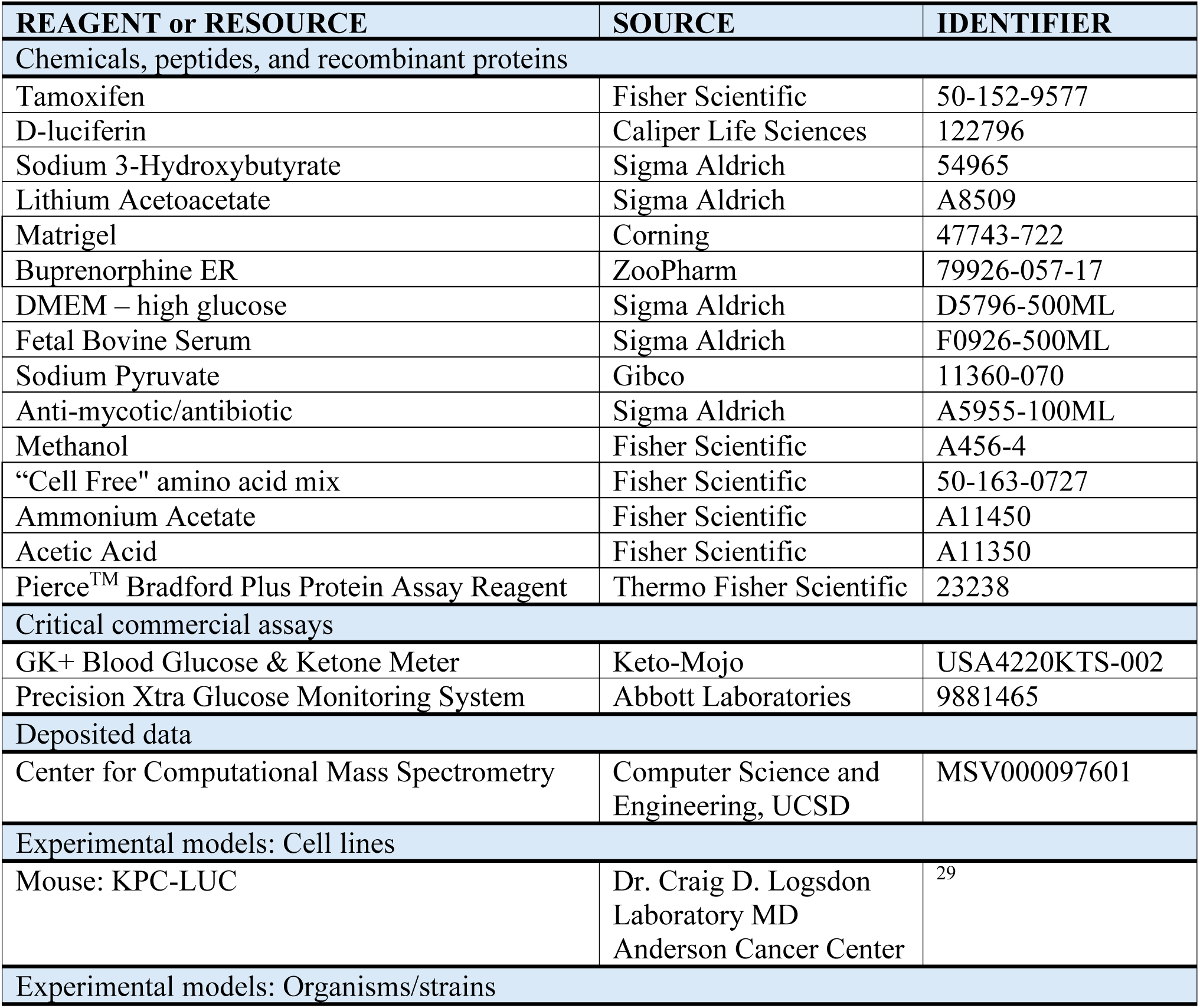
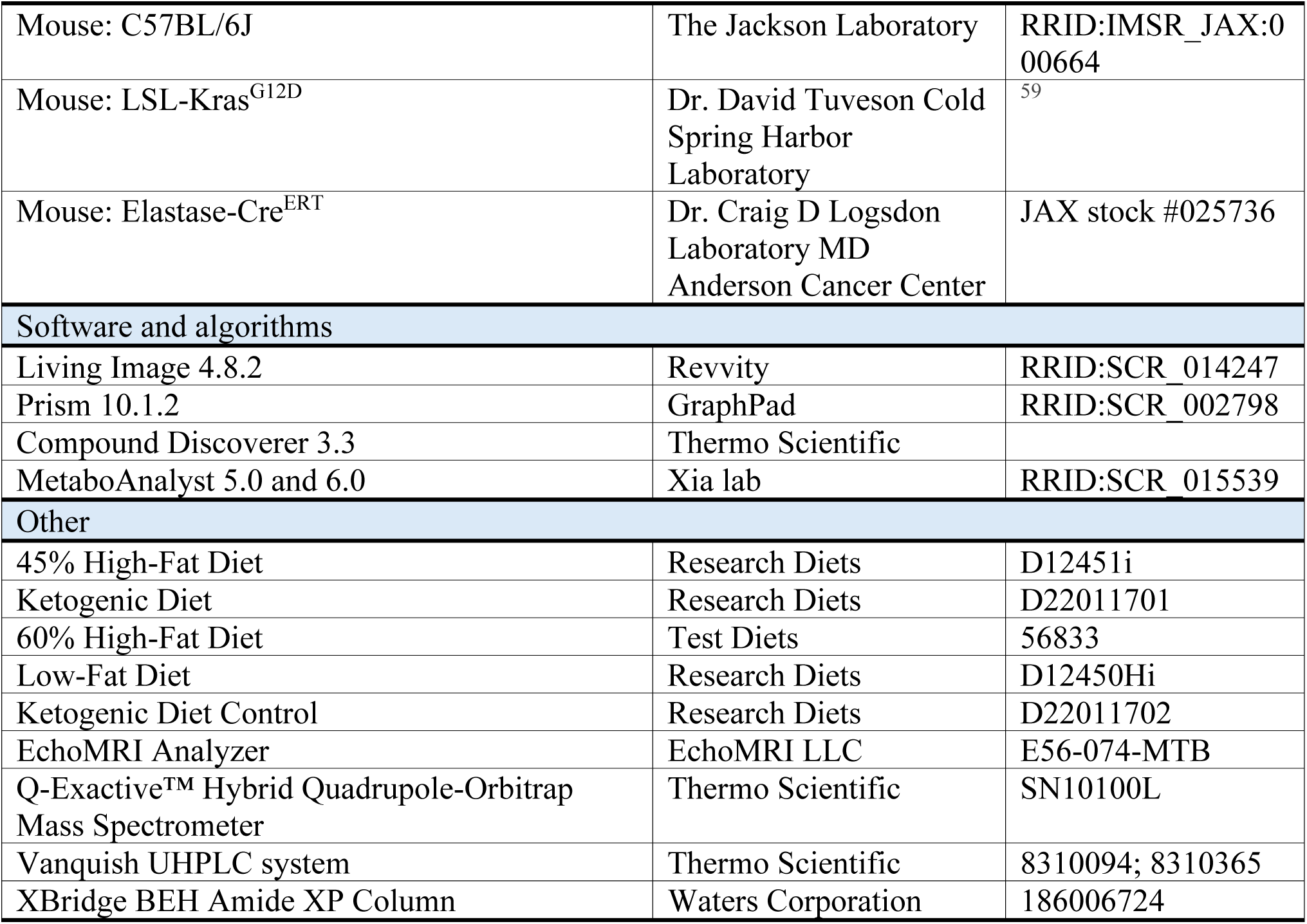

### Experimental model details

#### PDAC genetically engineered and orthotopic mouse models

All animal studies were conducted in accordance with the Animal Research: Reporting of *in vivo* Experiments guidelines and followed all regulatory and ethical standards. The protocols were reviewed and approved by the Institutional Animal Care and Use Committee at The Ohio State University, ensuring compliance with institutional and national guidelines for the ethical treatment of animals in research.

LSL-Kras^G12D^ mice^59^ were bred with Elastase-CreErT mice^60^ to make Kras^G12D/^CRE mice (mixed background) and littermates (control) and at 40 days old were administered tamoxifen orally (3mg/40g body weight) for four consecutive days, as previously described.^32^ Body weight was measured weekly.

For the orthotopic models, pancreatic tumor cells derived from a KPC GEMM expressing firefly luciferase (KPC-LUC) were utilized in the PDAC orthotopic model.^29^ Viable cells were counted and mixed in phosphate-buffered saline with 20% Matrigel (Corning). C57BL/6J male mice (The Jackson Laboratories) were given buprenorphine extended release (ZooPharm) for pain relief and then orthotopically injected into the pancreas with KPC-LUC cells (0.25 x 10^6^ cells/mouse) as previously described.^29^ Subcutaneous injections of D-luciferin (1.5 mg/mouse; Caliper Life Sciences) were used to visualize tumor growth weekly using the In Vivo Imaging System (IVIS; Caliper Life Sciences) and analyzed using the Living Image software.^29^

#### Method details Cell culturing

Cell lines were cultured at 37°C with 5% CO_2_ in DMEM supplemented with 10% FBS, 1% sodium pyruvate, and 1% anti-mycotic/antibiotic.

#### Ketone body treatments

Murine KPC cells (500,000 cells per well) were plated in a 6-well plate and allowed to adhere overnight. The cells were then treated with either 5 mM NaHB (Sigma-Aldrich) or 5 mM LiAcAc (Sigma-Aldrich) in serum-free media for 48 hours. Vehicle control KPC cells were maintained in serum-free media for the same duration.

#### Diet interventions

Diet-induced obesity (DIO) was achieved using a high-fat diet (HFD) in which 45% or 60% of the energy was derived from fat (D12451i; Research Diets and 56833; Test Diets). Mice were fed a low-fat (non-obese) diet containing 10% energy derived from fat (D12450Hi; Research Diets) to maintain a non-obese state. After 15 weeks on either DIO or non-obese conditions, mice were either maintained on their respective diets or switched to a KD providing 90% energy derived from fat (D22011701; Research Diets) or a LFD, used as a KD control (KDC) (D22011702; Research Diets) where 12% of energy was derived from fat, for 6 weeks (**Supplemental Table 1**). Mice were fed *ad libitum* and food changed weekly, except for the KD mice where it was changed every two to three days per week.

#### Circulating ketone measurement

β-HB concentration was measured using GK+ Blood Glucose & Ketone Meter (Keto-Mojo) according to the manufacturer’s instructions using a tail prick blood sample. Measurements were done twice a week and averaged after the KD intervention to account for variations over time.

#### Glucose tolerance test

GTTs were performed after dietary interventions and prior to the end of study. Mice were fasted for 12 h before the start of GTT, and body weights after food deprivation were used to calculate glucose doses (1.0 g glucose/kg BW). Blood was collected via tail prick before glucose injection for the baseline measurement and at 15, 30, 45, 60, 90, and 120-min after glucose challenge. Glucose was measured with a Precision Xtra Glucose Monitoring System (Abbott Laboratories, Abbott, IL).

#### Body Composition

Body composition (percent lean and fat mass) was measured using an EchoMRI analyzer (EchoMRI LLC).

#### Histopathology and pancreas scoring

Histopathological analysis was performed on formalin-fixed, paraffin-embedded pancreatic tissue sections stained with hematoxylin and eosin (H&E). In addition, sections stained with Masson’s trichrome were evaluated for fibrosis. Sections were evaluated and scored by a board-certified veterinary pathologist (S.E.K.) blinded to genotype and intervention. Mouse pancreatic intraepithelial neoplasia lesions were classified and graded according to standard criteria.^61–63^ Pancreata were also scored for the most severe and frequent lesions per section using a scoring scheme adapted from a grading scheme used for lesions in transgenic adenocarcinoma of the mouse prostate.^64^ Fibrosis and inflammation were assessed on a scale of 0 to 3. The adjusted lesion scores were added to the fibrosis and inflammation scores to generate the total pathological score (**Supplemental Table 2**).

#### Mass spectrometry-based metabolomics

Intracellular metabolites from each biological replicate were extracted using a cold methanol-based extraction. Cells were harvested, counted, and 1 × 10^6^ cells were aliquoted in triplicate for metabolite extraction. Cells were washed using cold PBS before the addition of 250 μL of LC-MS grade methanol (Fisher Scientific). For metabolite extraction of pancreas tissue samples, 30µg of tissue was homogenized and incubated with 250µL of 4:1 (v/v) methanol and water. Internal standards containing ^13^C and ^15^N labeled amino acids mix (1.2 mg/mL methanol) were added to the samples at a volume of 50 μL (Cambridge Isotope Laboratories).

Metabolomics samples were homogenized for 2 min before incubating at −20 °C for 20 min to extract polar metabolites. Samples were pelleted, and 200 μL of supernatant was transferred to an LC-MS vial for analysis. A pooled quality control (pQC) sample was collected by aliquoting an equal volume of supernatants from all biological samples into a separate vial and homogenized by vortexing. Sample vials were maintained in the autosampler tray at 4 °C.

LC-MS/MS analysis of pancreas tissue samples was conducted using a Vanquish UHPLC system (Thermo Scientific, Waltham, MA, USA) coupled to a Q-Exactive™ Hybrid Quadrupole-Orbitrap Mass Spectrometer (Thermo Scientific, Waltham, MA, USA). A 5 µL sample was injected onto an XBridge BEH Amide XP Column (150 mm × 2.1 mm, 130 Å, 2.5 µm particle size) (Waters Corporation, Milford, MA, USA) with the column temperature maintained at 40°C.

The mobile phases used were: Phase A, consisting of 5 mM ammonium acetate (Fisher Scientific) in acetonitrile/water (10:90, v/v) with 0.1% acetic acid (Fisher Scientific), and Phase B, which included 5 mM ammonium acetate in acetonitrile/water (90:10, v/v) with 0.1% acetic acid. The flow rate was set at 0.3 mL/min for a 20-minute acquisition, following a stepwise gradient for solvent B: initially 70%; from 0–5 min, 30%; from 5–9 min, 30%; and from 9–11 min, 70%. A divert valve was engaged to direct the flow to waste during the final 0.01 minutes and the first 0.5 minutes of the run.

The Q-Exactive™ system was equipped with an electrospray ionization (ESI) source, operated in both negative and positive ion modes, optimizing the detection of a wide range of metabolites. In positive mode, the MS spectra were collected in Full MS-SIM mode at a resolution of 70,000, AGC target of 3×10^6^, maximum IT of 200 ms, chromatographic peak width of 6 seconds, and scan range of 60-900 m/z. The tune file for positive detection used a sheath gas flow rate of 50, auxiliary gas flow rate of 15, sweep gas flow rate of 1, spray voltage of 3.5 kV, capillary temperature of 350 °C, S-lens RF level of 50.0, and auxiliary gas temperature of 425°C. Negative mode used the same parameters except for a spray voltage of −2.75 kV.

MS-based untargeted metabolomics was performed in triplicates. pQC samples were included at the beginning and end of the run, with a pQC sample followed by a solvent blank inserted after every ten biological sample injections. pQC samples were also used for data dependent acquisition (DDA) of the top 10 most abundant ions with dynamic exclusion for compound identification. All raw data from the MS analysis has been deposited in MassIVE spectral database under accession number MSV000097601.

All raw UHPLC−MS spectral data were imported into the feature annotation and interpretation software Compound Discoverer 3.3 (Thermo Scientific, Waltham MA, USA) for metabolite identification. The MS data were queried against an in-house database containing experimentally obtained MS/MS spectra of authentic analytical standards with mzCloud database and Compound Discoverer to 3.3 for metabolite identification. The collected area counts were then normalized to cell count per replicate or protein content of biological samples, and spectra were filtered to reduce redundancy and ensure instrument reproducibility. Protein levels of individual samples were quantified using Pierce^TM^ Bradford Plus Protein Assay Reagent (Fisher Scientific, Hampton, NH), using manufacturer recommended protocols. Any metabolite with a coefficient of variance more significant than 20% was removed before further analysis.

Statistical analyses, including univariate (T-test), multivariate (PLS-DA) and pathway analysis, were conducted using the online resources MetaboAnalyst 5.0 (PDAC cells) and MetaboAnalyst 6.0 (pancreas tissue of PDAC GEMMs) using the statistical analysis and enrichment analysis modules. PLS-DA was used for the interpretation of the metabolic differences between the ketone body treatments and pancreas tissue from the obese PDAC GEMMs subjected to dietary interventions. The global test package was used to examine metabolites within respective pathways to determine their association with an experimental variable. Primary metabolites from top enriched pathways were selected for further analysis.

#### Statistical analysis

Statistical analyses were conducted using Prism 10.1.2 (GraphPad Software, San Diego, CA). To compare one variable between two groups, unpaired t-tests were used, or Mann-Whitney tests were used if the data were not normally distributed. For assessing differences over time between groups, repeated measures mixed-between-within ANOVA was used. Comparisons involving more than two groups were analyzed using one-way ANOVA, a one-way ANOVA with Brown-Forsythe correction for unequal variance, or the Kruskal-Wallis test for non-normally distributed data. Tukey’s, Dunnett’s or Dunn’s correction for multiple testing was used, respectively. Two-way ANOVA was utilized to examine interactions between multiple factors and Tukey’s multiple testing correction was used for multiple comparisons. Results are presented as the mean ± SD unless otherwise specified in the figure legends. Statistical significance was defined as *p≤0.05, **p≤0.01, ***p≤0.001, and ****p≤0.0001.

## Acknowledgments

We thank The Ohio State University Comprehensive Cancer Center Small Animal Imaging Core (SAIC) Laboratory for their IVIS and EchoMRI services, Comparative Pathology and Digital Imaging Shared Resource (CPDISR) for IHC support and histopathology evaluation. Research reported in this publication was partially supported by Start-up funds from The Ohio State University Comprehensive Cancer Center and The National Cancer Institute R01CA279707 (Z. Cruz-Monserrate), Pelotonia Scholarship Program (V Pita-Grisanti and G Vulic), The National Cancer Institute T32 Tumor Immunology Fellowship T32CA09223 (K Chasser), the GR111977 USDA Training grant (E. Velez-Bonet) and the OSUCCC P30 CA016058 National Cancer Institute. The content in this article is solely the responsibility of the authors and does not necessarily represent the official views of the National Institutes of Health. Any opinions, findings, and conclusions expressed in this material are those of the author(s) and do not necessarily reflect those of the Pelotonia Scholarship Program, The Ohio State University, or the National Institutes of Health. Overall experimental design figures created with Biorender.com

## Author contributions

E.V-B. and Z.C-M. conceived the study. E.V-B. designed and performed most of the experiments. K.C., Z.H., H.Y.H, V.P.G., A.L., G.V., K.G-F., F.C., H.Z., Z.J., J.S.V., S.E.K. contributed to the investigation. E.V-B, K.C., K.G-F, F.C., H.Z., Z.J., S.E.K., S.C. and Z.C-M conducted formal analysis. F.C., H.Z., Z.J., and J.S.V. also contributed resources. E.V-B, K.G-F, and Z.C-M wrote and revised the manuscript. E.V-B, K.G-F, and Z.C-M contributed to the discussion and critical review of the manuscript. Z.C-M. supervised and funded the project. All authors reviewed the final manuscript.

