## Supplemental Tables and Figures for "Ketogenic diet prevents obesity-associated pancreatic cancer independent of weight loss and induces pancreatic metabolic reprogramming"

| Product # | D12451i | D22011701 | D22011702 | 56833 | D12450Hi |
|---|---|---|---|---|---|
| | High-Fat Diet **DIO (45%)** | Ketogenic Diet **KD** | Ketogenic Diet Control **KDC** | High-Fat Diet **DIO (60%)** | Low-Fat Diet **non-obese** |
| **Macronutrient Energy Distribution (%kcal)** | | | | | |
| Protein | 20 | 9 | 19 | 18 | 20 |
| Carbohydrate | 35 | 1 | 70 | 20 | 70 |
| Fat | 45 | 90 | 12 | 62 | 10 |
| **Ingredients (gm)** | | | | | |
| Corn Oil | 0 | 55 | 50 | 0 | 0 |
| Soybean Oil | 25 | 0 | 0 | 32.31 | 25 |
| Lard | 177.5 | 325 | 0 | 316.6 | 20 |
| Casein | 200 | 96.5 | 200 | 258.45 | 200 |
| DL-Methionine | 0 | 1.25 | 300 | 0 | 0 |
| Corn Starch | 72.8 | 0 | 500 | 0 | 452.2 |
| Maltodextrin | 0 | 0 | 150 | 161.53 | 0 |
| Sucrose | 176.8 | 0 | 0 | 88.47 | 176.8 |
| Cellulose | 0 | 50 | 50 | 0 | 0 |
| Milk Fat, Anhydrous | 0 | 0 | 0 | 0 | 0 |
| Mineral Mix S10001A (without Ca & P) | 0 | 13.1 | 13.1 | 0 | 0 |
| Mineral Mix S10026B | 50 | 0 | 0 | 0 | 50 |
| DIO Mineral Mix | 0 | 0 | 0 | 12.92 | 0 |
| Calcium Phosphate, Dibasic | 0 | 14.3 | 14.3 | 0 | 0 |
| Calcium Phosphate, Monobasic | 0 | 5 | 5 | 0 | 0 |
| Calcium Phosphate | 0 | 0 | 0 | 16.8 | 0 |
| Vitamin Mix V10001 | 0 | 9.8 | 9.8 | 0 | 0 |
| Vitamin Mix V10001C | 1 | 0 | 0 | 0 | 1 |
| AIN-76A Vitamin Mix | 0 | 0 | 0 | 12.92 | 0 |
| Choline Bitartrate | 2 | 2 | 2 | 2.58 | 2 |
| Cystine, L | 3 | 0 | 0 | 3.88 | 3 |
| Lodex 10 (Maltodextrin) | 100 | 0 | 0 | 0 | 75 |
| Fiber | 50 | 0 | 0 | 0 | 50 |

**Supplemental Table 1**: **Composition of diets.** The table summarizes the relative contribution of protein, carbohydrates, and fat to total caloric content across five distinct rodent diets formulated for metabolic research and gram quantities of individual ingredients per diet are listed.

| Lesion Grade | Diagnosis | Distribution | Adjusted Lesion Score |
|---|---|---|---|
| 0 | Normal | Diffuse | 0 |
| 1 | Low-grade (mPanIN-1, 2) | Focal (F) | 1 |
| 1 | Low-grade (mPanIN-1, 2) | Multifocal (MF) | 2 |
| 1 | Low-grade (mPanIN-1, 2) | Diffuse | 3 |
| 2 | High- grade (mPanIN-3) | Focal (F) | 4 |
| 2 | High-grade (mPanIN-3) | Multifocal (MF) | 5 |
| 2 | High-grade (mPanIN-3) | Diffuse | 6 |
| 3 | PDAC | Focal (F) | 7 |
| 3 | PDAC | Multifocal (MF) | 8 |
| 3 | PDAC | Diffuse | 9 |
| **Fibrosis Scoring** | | | |
| 0 | Normal and no Fibrosis | | |
| 1 | 1-10% increase in collagenous stroma and occasional; periductular or scant perilobular fibrosis | | |
| 2 | 10-30% increase in collagenous stroma showing moderate fibrosis | | |
| 3 | 30-50% increase in collagenous stroma showing extensive fibrosis | | |
| 4 | More than 50% fibrosis | | |
| **Inflammation Scoring** | | | |
| 0 | No inflammation | | |
| 1 | Minimal infiltration of periductal tissue and occasional scattered leukocytes | | |
| 2 | Inflammation around ducts and extending into the parenchyma (< 50% of lobules) with moderate numbers of leukocytes | | |
| 3 | Inflammation around ducts and extending into the parenchyma (51%-75% of lobules) with dense aggregates of leukocytes | | |
| 4 | Inflammation around ducts and extending into the parenchyma (>75% lobules) | | |

**Supplemental Table 2: Scoring system for adjusted pancreatic lesions, fibrosis, and inflammation.** An adjusted lesion scoring system evaluates both the most severe and most common pancreatic lesions on a scale of 0-3 with an adjusted score based on their distribution (focal, multifocal, or diffuse; 0-9) resulting in two adjusted scores. Distributions of these lesions are determined and described as focal if less than 3 foci contain the lesion, multifocal if there are three or more foci or less than 50% of the section containing the lesion or diffuse if greater than 50% of section contains the lesion. The adjusted scores are then added to obtain a sum. The adjusted lesion score sum is added to the fibrosis and inflammation scores to generate the total pathological score.

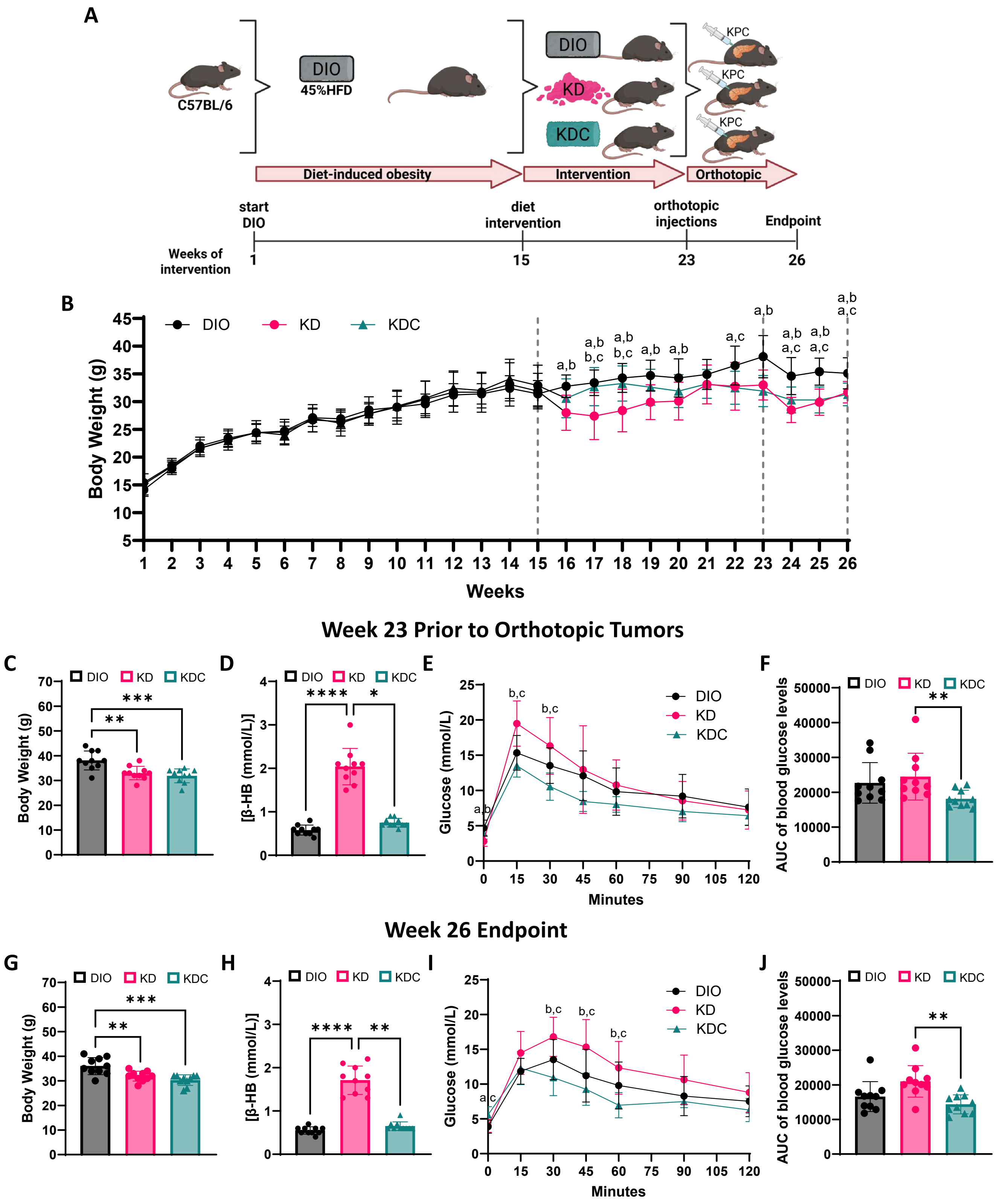


**Supplemental Figure 1**

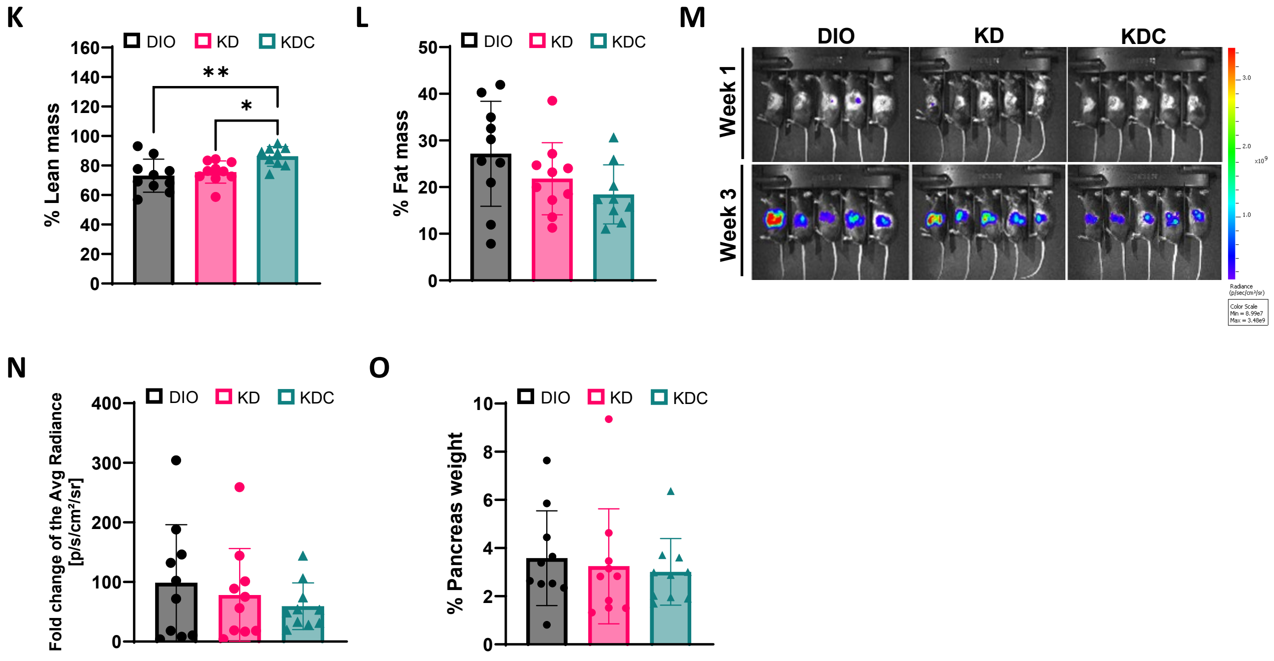


**Supplemental Figure 1: KD does not inhibit tumor growth but induces weight loss in an obese orthotopic model of PDAC.**

**A**) Experimental design to evaluate the effects of the KD and KDC following DIO in an orthotopic PDAC model using a 45% HFD. **B**) Body weight of C57BL/6J mice over 26 weeks (n=30). **C**) Body weight prior to orthotopic tumors (week 23) after randomization to KD and KDC (n=10 mice/group). **D**) β-HB concentration prior to orthotopic tumor implantation. GTT measuring **E**) glucose and **F**) AUC prior to orthotopic tumor implantation. **G**) Body weight at endpoint (week 26). **H**) β-HB concentration at endpoint. GTT measuring **I**) glucose and **J**) AUC at endpoint. Percent **K**) lean mass and **L**) fat mass of mice at endpoint. **M**) Representative bioluminescence imaging used to measure tumor growth. **N**) Fold change of the average tumor radiance at endpoint. **O**) Percent pancreas weight. Statistical significance determined by mixed-between-within ANOVA's where significance between groups indicated by letter pairs listed for the diet groups: a) DIO, b) KD and c) KDC (**B**, **E**, and **I**), one-way ANOVA with Tukey's multiple comparisons (**C**, **G**, **K**, and **L**), or Kruskal-Wallis test with Dunn's correction for multiple testing (**D**, **F**, **H**, **J**, **N**, **and O**). *p<0.05, **p<0.01, ***p<0.001, ****p<0.0001

**Supplemental Figure 1-continued**

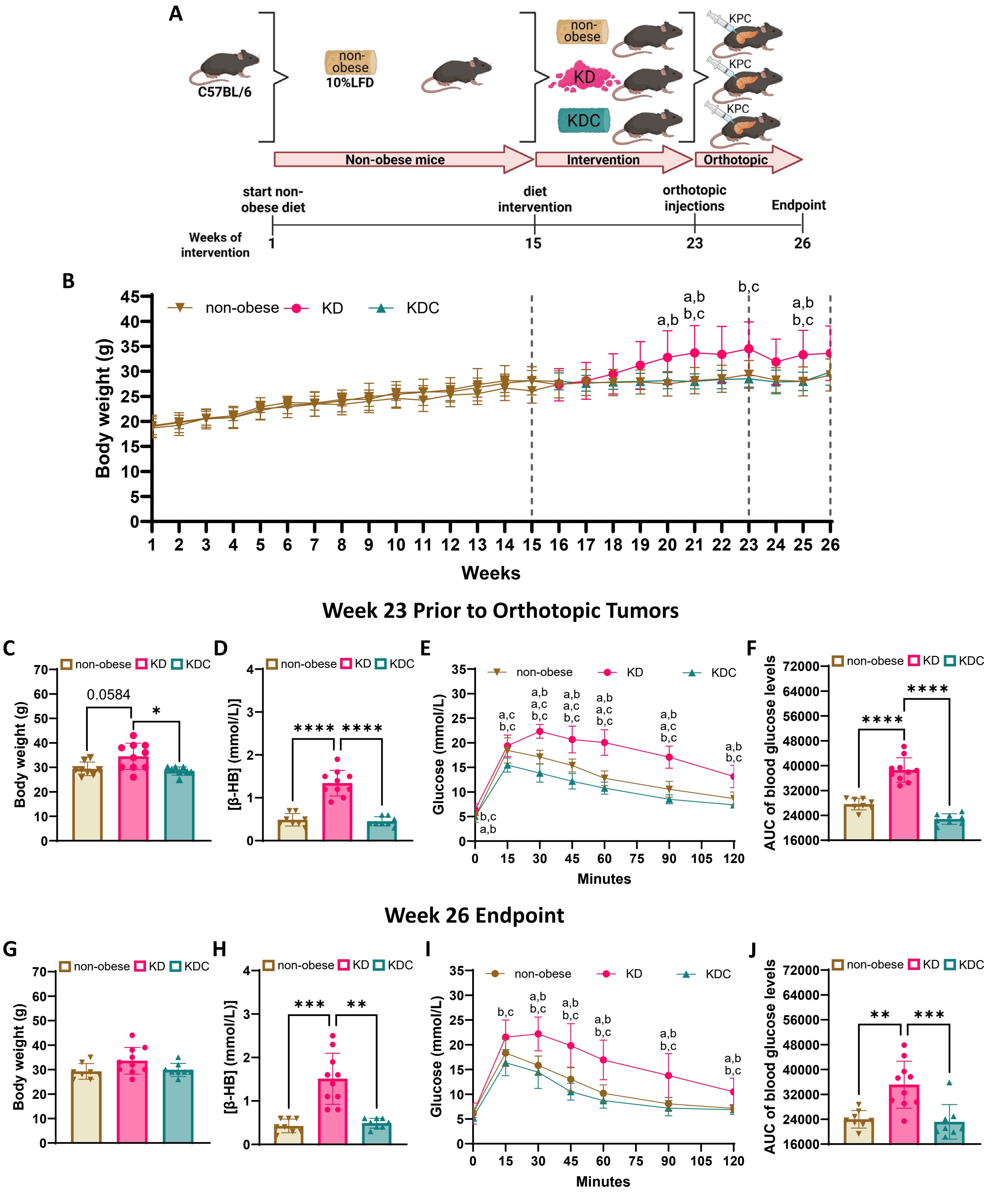


### Supplemental Figure 2

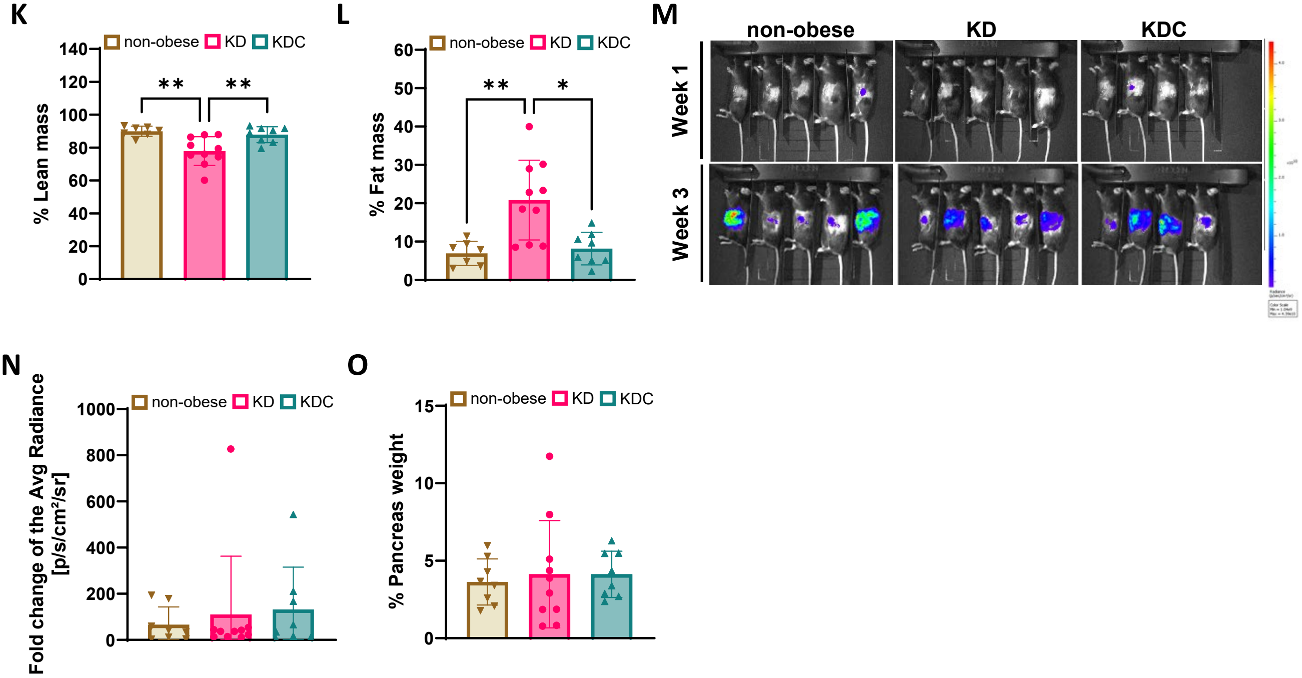


**Supplemental Figure 2: KD does not inhibit tumor growth in a non-obese orthotopic model of PDAC.**

**A**) Experimental design to evaluate the effects of the KD and KDC in an orthotopic PDAC model using a 10% low-fat diet. **B**) Body weight of C57BL/6J mice over 26 weeks (n=30). **C**) Body weight prior to orthotopic tumors (week 23) after randomization to KD and KDC (n=8-10 mice/group). **D**) β-HB concentration prior to orthotopic tumor implantation. GTT measuring **E**) glucose and **F**) AUC prior to orthotopic tumor implantation. **G**) Body weight at endpoint (week 26). **H**) β-HB concentration at endpoint. GTT measuring **I**) glucose and **J**) AUC at endpoint. Percent **K**) lean mass and **L**) fat mass at endpoint. **M**) Representative bioluminescence imaging used to measure tumor growth. **N**) Fold change of the average tumor radiance at endpoint. **O**) Percent pancreas weight. Statistical significance determined by mixed-between-within ANOVA's where significance between groups indicated by letter pairs listed for the diet groups: a) non-obese, b) KD and c) KDC (**B**, **E**, and **I**), one-way ANOVA with Tukey's multiple comparisons (**C**, **G**, and **K**), one-way ANOVA with Brown Forsythe and Dunnet's corrections (**L**) or Kruskal-Wallis test with Dunn's correction for multiple testing (**D**, **F**, **H**, **J**, **N**, **and O**). $^*p<0.05$, $^{**}p<0.01$, $^{***}p<0.001$, $^{****}p<0.0001$.

### Supplemental Figure 2 (cont.)
